# Intravenous lipid-siRNA conjugate mediates gene silencing at the blood-brain barrier and blood-CSF barrier

**DOI:** 10.1101/2025.03.14.642142

**Authors:** Alexander G. Sorets, Katrina R. Schwensen, Nora Francini, Andrew Kjar, Sarah Lyons, Joshua C. Park, Dillon Palmer, Adam M. Abdulrahman, Rebecca P. Cowell, Ketaki A. Katdare, Ella N. Hoogenboezem, Angela Wang, Neil Dani, Craig L. Duvall, Ethan S. Lippmann

## Abstract

Barriers of the central nervous system (CNS), such as the blood-brain barrier (BBB) and blood-cerebrospinal fluid barrier (BCSFB), regulate the two-way exchange of material between the blood and CNS. These barriers pose a considerable challenge for efficacious delivery of intravenously administered therapies into the CNS, motivating exploration of their function and ways to modulate their properties. While the BBB and BCSFB can become dysfunctional in patients with chronic CNS diseases, few studies have focused on strategies for targeting these interfaces. Here, we showed that an intravenously administered albumin-binding lipid-siRNA conjugate was delivered to and silences genes within brain endothelial cells and choroid plexus epithelial cells, which comprise the BBB and BCSFB, respectively. A single intravenous dose of lipid-siRNA conjugate was delivered to ∼100% of brain endothelial cells and major choroid plexus cell types, without any substantial delivery into brain parenchymal tissue. Sustained gene silencing was achieved in both brain endothelial cells (over two weeks) and bulk choroid plexus tissues (up to one month). Moreover, single cell RNA sequencing demonstrated gene knockdown in capillaries, venous endothelial cells, and choroid plexus epithelial cells without silencing genes in parenchymal cell populations. Collectively, this work establishes an effective nonviral framework to mediate gene inhibition in the brain barriers.

**GRAPHICAL ABSTRACT:** 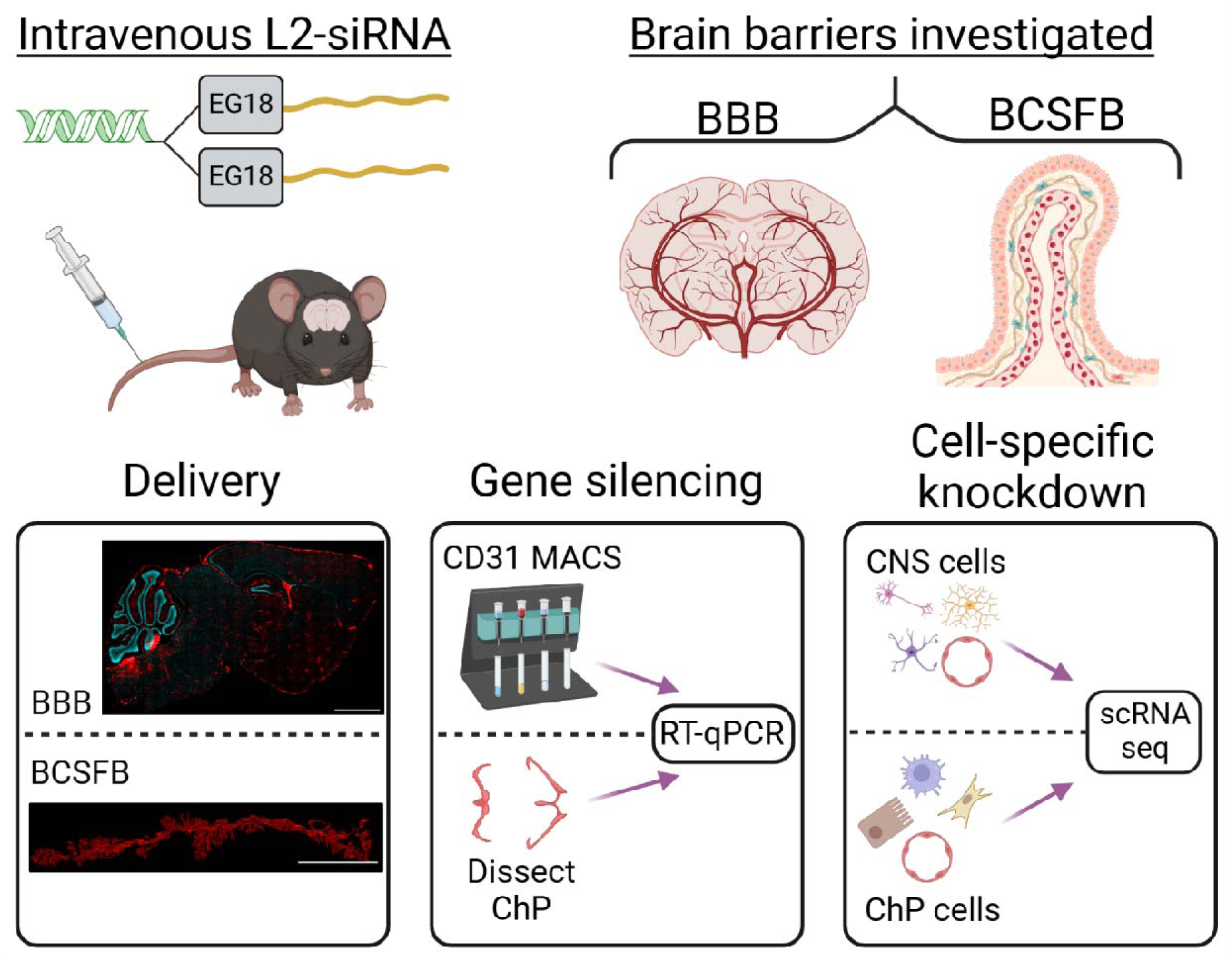

## INTRODUCTION

Blood-central nervous system (CNS) barriers pose a formidable obstacle to achieving drug delivery to the brain. These interfaces include the blood-brain barrier (BBB), which describes the restrictive properties of brain endothelial cells, as well as the blood-cerebrospinal fluid (CSF) barrier (BCSFB) formed by the epithelial cells in the choroid plexus [1]. By limiting non-specific transport, the BBB and BCSFB collectively mediate selective exchange of materials between the CNS and the rest of the body, including nutrients, cellular waste products, and immune cells [2]. Yet, these homeostatic functions of the blood-CNS barriers are perturbed in many pathological conditions, exacerbating or even initiating disease progression [3–6]. Targeting cells of the brain barriers is an underexplored avenue for modulating barrier function as a direct disease treatment or as an approach to potentiate delivery of a secondary therapy for treating diseases within the CNS.

The vasculature serves as the principal component of the BBB and may be an important target for drug delivery. Compared to other endothelial cells, brain endothelial cells possess unique features, such as a continuous tight junction network that prevents transport between cells and a mechanism to suppress caveolae-mediated transcytosis through cells [7–9]. BBB dysfunction, characterized by loss of these properties, occurs in a number of diseases including Multiple Sclerosis, Alzheimer’s Disease, and Huntington’s disease [10,11]. These changes may cause alteration in normal local ion concentrations, increased plasma protein entry, or immune cell infiltration, collectively leading to neuronal degeneration [9]. To mitigate this damage, several viral vectors have been engineered to enhance brain endothelial-specific tropism in rodents. Notable examples include AAV-PHP.V1 [12], AAV-BR1 [13], AAVBI30 [14], and AAV.X1 [15], which have been used to answer research questions about cerebrovasculature as well as rescue of neurological dysfunction. While AAVs are an effective tool for modulating gene expression in the BBB, several hurdles may hinder their translational potential, such as the lack of suitability for repeated administration with some constructs, the potential for an immune response, and the limited transgene size capacity of some viruses [16]. Due largely to the safety and cost challenges of viral therapies, there is a need for non-viral therapeutic platforms that can safely modulate gene expression in vasculature that comprise the BBB.

In addition, few clinically-relevant techniques have been developed to target the choroid plexus (ChP), specifically the choroid plexus epithelial cells (CPECs) which constitute the functional unit of the BCSFB [17]. Despite comprising ∼1% of the brain weight, the choroid plexus forms ∼50% of the cerebral capillary surface area, highlighting its remarkable potential for transport [18,19]. In contrast to brain endothelial cells, ChP blood vessels are fenestrated, permitting leakage of blood-borne molecules into the stromal space which is inhabited by macrophages and fibroblasts. CPECs form the BCSFB by partitioning CSF from blood through a continuous network of tight junctions. CPECs thereby generate CSF as an ultrafiltrate of blood, driving CSF flow through the lateral, third, and fourth ventricles before filling the subarachnoid space surrounding the brain. Proper production of CSF by the ChPs is essential for development and maintenance of the CNS [20], as CPEC dysfunction plays a role in edema and hydrocephalus [21]. Further, because it functions as a conduit for immune cell entry, CPEC activity is inextricably linked to neuroinflammation [22]. Similar to methods used to target brain endothelial cells, viral approaches have emerged as a powerful tool to overexpress or inhibit genes in CPECs. In particular, the AAV2/5 serotype has been shown to sustain transduction in neonatal mice after intravenous delivery [23], but intracerebroventricular delivery is required for transduction but in adult mice [17,24]. Thus, at present, AAVs have not yet demonstrated efficacy from systemic administration, and new approaches are needed to target ChP dysfunction.

While viral vectors have been extensively described in the literature for CNS barrier targeting, nonviral strategies are less well developed. Systemic delivery of synthetic oligonucleotides for gene manipulation is gaining considerable interest due to advances in nucleic acid chemistry, which endow resistance to degradation in the bloodstream. Yet, nucleic acids exhibit poor drug delivery properties, as they lack effective binding to proteins and cells, leading to a short circulation half-life and poor tissue penetration [25]. Oligonucleotide encapsulation into nanoparticles and conjugation to hydrophobic or targeting moieties have been used to improve deliver to various organs and tissues, including the brain [26–29]. While some LNPs can deliver mRNA to the BBB [30], very few strategies have focused on siRNA delivery to brain barriers as the end goal.

Here, we sought to explore intravenous delivery of an albumin-binding short interfering RNA (siRNA) as an approach for delivery to and gene silencing within brain barriers. This approach was motivated by the well-described behavior of the albumin-binding dye Evans blue. Evans blue is often used as a research tool to assess and visualize BBB and BCSFB leakage, but in normal physiological conditions Evans blue labels CNS blood vessels and the choroid plexus [31]. We previously developed a lipid-siRNA conjugate (termed L2-siRNA) that reversibly binds albumin in blood, leading to broad silencing in tumors and injured knee joint tissues [32,33]. In this manuscript, we assessed intravenous L2-siRNA delivery to brain endothelial cells and all major choroid plexus structures. A time course of gene silencing kinetics was established for brain endothelial cells, non-endothelial parenchymal cells, and choroid plexus. Single cell RNA sequencing (scRNA-seq) methodology was leveraged to determine cell-specific gene silencing in the brain barriers and parenchyma. Overall, we characterize a platform for mediating gene silencing in brain barriers, both as a tool for mechanistic studies and as an avenue for clinical intervention.

## MATERIALS AND METHODS

### Synthesis of oligonucleotides

Oligonucleotides were synthesized using standard solid-phase phosphoramidite chemistry as previously described [34]. In brief, nucleic acid strands were grown off of universal support-controlled pore glass columns on a MerMade synthesizer (BioAutomation). Cy5-labelled oligonucleotides were synthesized on Cy5-functionalized 1000Å CPG (Glen Research). Amidites were dissolved in acetonitrile (0.1M), except 2′-OMe uridine which requires 20% dimethylformamide additionally. For synthesizing L2-siRNA, the stearyl amidite is dissolved in dichloromethane:acetonitrile (3:1, v:v). Ancillary reagents include the activator (5-Ethylthio-1H-Tetrazole, 0.25 M in acetonitrile), CAP A (Tetrahydrofuran/ 2,6-Lutidine/ Acetic Anhydride), CAP B (20% acetic anhydride, 30% 2,6-lutidine in ACN), 3% Dichloroacetic acid (for detritylation, in DCM). Capping and oxidation was performed using 0.02M iodine in Tetrahydrofuran/ Pyridine/Water (70:20:10) and sulfurization to create a phosphorothioate bond was performed with 0.05M Sulfurizing Reagent II in Pyridine/Acetonitrile. For all antisense strands, 5′-(E)-Vinylphosphonate (VP) was added at the 5’ end using POM-vinyl phosphonate 2’-OMe-uridine CE-phosphoroamidite (LGC genomics). For unconjugated siRNA (no lipid), the sense strands were synthesized DMT-on for further purification. Sequences for all oligonucleotides can be found in Supplemental Table 1.

### Cleavage and deprotection of oligonucleotides

Sense strands were cleaved and deprotected in 28%–30% ammonium hydroxide plus 40% methylamine (2 hours, room temperature), while Cy5-labelled oligonucleotides were cleaved solely in 28%–30% ammonium hydroxide (room temperature, 20 hours). The antisense strands were cleaved and deprotected in 3% diethylamine solution containing 28%–30% ammonium hydroxide (20 hours, 35°C) according to standardized protocols [35].

### Purification and verification of oligonucleotides

Organic solvents were removed after cleavage and deprotection with a SpeedVac (Savant SpeedVac SPD 120 Vacuum Concentrator, ThermoFisher). After resuspending in water containing 5% methanol, the lipidated sense strands were purified by HPLC (Waters 1525 EF) using a Clarity Oligo-RP column (Phenomenex) under a linear gradient [60% mobile phase A (50 mM triethylammonium acetate in water) to 90% mobile phase B (methanol)].

Non-lipidated sense strands (DMT-on) were desalted using Gel-Pak cartridge (Glen Research) followed by purification under a linear gradient (85% to 40% mobile phase A). Full-length oligonucleotide fractions were dried with a SpeedVac. Next, the DMT protecting group was removed using 20% acetic acid (1 hour, room temperature) followed by desalting, sterile filtration, and lyophilization.

Antisense strands were purified using anion-exchange chromatography (10×150 mm Source 15Q, Cytiva). Mobile phase A contained 10 mM sodium acetate in 20% acetonitrile and mobile phase B contained of sodium perchlorate (0.6 M) also in 20% acetonitrile. A gradient was obtained as follows: 85% to 73% mobile phase A for 10 minutes, and to 60% mobile phase B for 14 minutes at a flow rate of 4 ml/min. Purified oligonucleotides were desalted using a HiPrep 26/10 desalting column (Cytiva), sterile filtered, and lyophilized.

Oligonucleotide mass was verified by Liquid Chromatography-Mass Spectrometry (LC-MS, ThermoFisher LTQ Orbitrap XL Linear Ion Trap Mass Spectrometer) over a C18 column (Waters XBridge Oligonucleotide BEH. A linear gradient was obtained as follows: 85% phase A (16.3 mM triethylamine – 400 mM hexafluoroisopropanol) to 90% phase B (methanol) over 10 minutes at 45°C.

### Animal husbandry

Adult C57BL/6J male mice from Jackson Laboratory (3-4-months old) were used in these studies. Mice were housed in Vanderbilt facilities under a 12-hour light/dark cycle with *ad libitum* access to food and water. All protocols were approved by the Institutional Animal Care and Use Committee (IACUC) at Vanderbilt University.

### Intravenous injections, euthanasia, and tissue collection

On the day of all *in vivo* studies, siRNA duplexes were annealed in 0.9% sterile saline by heating to 95°C and gradually cooling to 4°C in an Eppendorf Thermal Cycler (EP6337000027). The siRNA was concentrated with a 3K Amicon Ultra spin filter (UFC500324) as needed. A 20 mg/kg dose of 0.4 mM siRNA was administered via tail-vein injection.

At the experimental endpoint, mice were euthanized either with an intraperitoneal injection of ketamine (450 mg/kg)/xylazine (50 mg/kg) or isoflurane overdose. Mice were then transcardially perfused with cold heparinized (10 U/ml) DPBS without calcium or magnesium (DPBS -/-). For flow cytometry, the brain was extracted, choroid plexuses were removed, and then the remaining brain tissue was processed as described below. For gene silencing studies, the choroid plexuses were manually dissected and stored in RNA Later (Thermo AM7020), and the remaining brain tissue was utilized for endothelial cell isolation. Biopsy punches (2 mm) were taken from major organs (liver, kidney, spleen, heart, lung) in consistent locations and stored in RNA Later. For biodistribution studies, mice were perfused for an additional 4 minutes with 4% paraformaldehyde (PFA) or 10% Formalin, and organs, brain, and spinal cord were immersion-fixed for an additional 24 hours at 4^◦^C.

### Whole brain dissociation

Extracted whole brains were dissociated into a single-cell suspension with the mouse adult brain dissociation kit (Miltenyi 130-107-677) according to manufacturer instructions. In brief, the brain was dissociated with a papain-based enzyme cocktail before the myelin was removed and the remaining red blood cells were lysed. To avoid contamination of non-barrier forming endothelial cells in final suspensions, the choroid plexuses were manually removed prior to dissociation when performing flow cytometry and bulk gene silencing analyses.

### Flow cytometry

After dissociation into a single-cell suspension, the cells were FcR blocked for 10 minutes on ice to prevent non-specific binding (10 µl per sample, Miltenyi 130-092-575). The cells were then stained with cell-specific markers for 30 minutes at 4°C: ACSA2 for astrocytes (1:2,000; Miltenyi 130-123-284), CD11b for microglia/macrophages (1:2,000; BD biosciences 561689), CD31 for endothelial cells (1:2,000; eBioscience 17-0311-80), and O1 for oligodendrocytes (1:100, R&D systems FAB1327G). After washing three times with DPBS-/- plus 0.5% BSA, cells were resuspended in the same buffer with DAPI (1:10,000, 5 mg/ml stock, Thermo D1306) and run on a BD LSR Fortessa flow cytometer. More than 70,000 live cell events were recorded for each experimental sample. Percent Cy5 positive cells was determined by gating off fluorescence-minus-one (FMO) controls originating from an uninjected mouse. Standard single-color compensation controls were run for each fluorophore. The gating scheme is shown in Supplemental Fig. 1.

### Magnetic activated cell sorting

CD31^pos^ endothelial cells were isolated from dissociated brains with anti-CD31 conjugated magnetic beads (Miltenyi Biotec 130-097-418) according to the manufacturer’s instructions. After incubation with beads, cells were sorted on a magnetic column into CD31^neg^ and CD31^pos^ populations. Each group was immediately processed for RT-qPCR as described below.

### RT-qPCR

RNA extractions were tailored for the specific sample type (cells isolated by MACS, choroid plexuses, organs). From whole organs, tissue samples were collected with biopsy punches and homogenized in RLT buffer plus 1:100 β-mercaptoethanol (RLT+) with 5 mm stainless steel beads (Qiagen cat. No. 69989) for 5 minutes at 30 Hz (TissueLyser II). Organ homogenates were then processed according to manufacturer instructions with the RNeasy Plus mini kit (Qiagen 74134). Sorted brain cell populations were lysed by resuspending in RLT+, and CD31^neg^ lysates were extracted with the RNeasy Plus mini kit, while CD31^pos^ lysates were extracted with the RNeasy Plus micro kit (Qiagen 74034). Each dissected choroid plexus (both laterals and 4^th^ ventricle) was extracted by homogenization in RLT + with 5 mm stainless steel beads for 5 minutes at 30 Hz. The choroid plexus homogenates were then processed with the RNeasy plus micro kit.

After the RNA was eluted into RNAse-free water, the concentration and purity were measured by 230/260/280 nm absorbance on a Nanodrop 2000c spectrophotometer. An equivalent mass of RNA was loaded into each cDNA reaction, which was performed according to iScript (BioRad 1708891) manufacturer instructions (5 min @ 25^◦^C, 20 @ 46^◦^C, 1 min @ 95^◦^C, hold @ 4^◦^C). qPCR was performed by preparing a 20 µl reaction mixture composed of 2X master mix (Thermo 4304437), water, Taqman probes, and cDNA sample. The following TaqMan probes were used: Mm00478295_m1 (*Ppib*), Mm01245874_g1 (*Rpl27*), Mm00727012_s1 (*Cldn5*), Mm01253033_m1 (*Gfap*), Mm00479862_g1 (*Iba1*), and Mm02619580_g1 (*Actb*). Samples were run in duplicate on a Biorad CFX96 and analyzed in CFX Maestro software using the ΔΔCt method. Choroid plexuses, isolated cells, and lungs were normalized to *Rpl27*, while all other organ samples were normalized to *Actb.* Conventional RT-qPCR controls (no-template control and no reverse transcriptase control) were run once for each probe/sample combination and did not amplify.

### Peptide nucleic acid (PNA) hybridization assay

As previously described, we used the PNA assay to quantify absolute siRNA delivery to organs [34,36]. Tissue biopsy punches were homogenized in 300 μl of buffer (Thermo QS0518) plus proteinase-K (Thermo QS0511, 1:100), and disrupted using a Tissuelyzer 2.0 for 5 minutes at 30 Hz. Supernatant was collected and stored after a 1-hour incubation at 65°C. Potassium chloride (3M) was added to precipitate SDS, and then samples were annealed to the Cy3-labeled PNA probe (∼10 pmol/ 150 ul of sample, PNA bio) in hybridization buffer (50 mM Tris, 10% ACN, pH 8.8). The annealed samples were run through liquid chromatography (Shimadzu) using a DNAPac PA100 anion-exchange column (Thermo Fisher Scientific). A gradient was established with mobile phase A (50% acetonitrile and 50% 25 mM Tris–HCl, pH 8.5; 1 mM ethylenediaminetetraacetate in water) and buffer B (800 mM sodium perchlorate in buffer A): 10% buffer B within 4 minutes, 50% buffer B for 1 minute and 50% to 100% buffer B within 5 minutes. The final mass of siRNA was calculated using the area under the curve of Cy3 fluorescence from a standard curve of known quantities of L2-siRNA spiked into untreated tissue homogenates. The siRNA quantity was normalized to mass of each tissue sample.

### Histology

To prepare fixed brains for frozen sections, sucrose gradients were performed for cryoprotection by immersion in 15% sucrose for 24 hours, followed by immersion in 30% sucrose for an additional day. The brains were then embedded in Epredia Neg-50 medium (Fisher 22-110-617), sectioned to 20 μm on a cryostat, and stored at -80°C until staining or stored in PBS and stained immediately as floating sections.

Antibody staining on frozen slides was performed as follows. First, samples were thawed to room temperature and a barrier was drawn using a hydrophobic pen to localize the staining reagents on the sample. Samples were washed in DPBS-/-followed by blocking for 1 hour at room temperature with 5% donkey serum (Sigma D9663). Slides are then incubated overnight at 4°C with the primary antibody diluted in 1% BSA in DPBS. Primary antibodies include CD31 (1:100, BD biosciences 550539), AQP4-488 (1:500, Abcam, ab284135), AQP1 (1:200, Abcam ab168387), and Iba1 (1:500, Wako 019-19741). The sections were then washed three times in DPBS for 5 minutes, followed by a 1-hour room temperature incubation in the appropriate secondary antibody (1:500 dilution in DPBS containing 1% BSA). Sections were washed, then incubated with DAPI (1:5,000-1:10,000, 5 mg/ml stock, Thermo D1306) for 10 minutes, washed again in DPBS, and then mounted under a coverslip with ProLong gold antifade reagent (ThermoFisher P36930). Imaging was performed either on a Leica epifluorescence microscope (tiled images of entire brain and organs) or a Zeiss LSM 710 confocal microscope. Image processing was performed using Fiji software. For samples where antibody staining was not required (i.e. assessing Cy5-tagged siRNA signal), slides were washed twice with DPBS and then mounted with ProLong gold plus DAPI.

### Quantification of Cy5-labeled L2-siRNA and siRNA in the choroid plexus

The mean fluorescence intensity of the Cy5 signal in the lateral, 3^rd^, and 4^th^ ventricle choroid plexus was quantified for each 20 μm section in ImageJ. The triangle thresholding method was applied to the nuclear stain (DAPI) channel to faithfully detect the edge of the tissue and bound the region. The tracing wand tool was then used to select the thresholded area, which was added to the ROI manager. Natural holes in the tissue were excluded with the boolean logic functions in the ROI manager. Once a complete ROI set had been created, it was applied to the Cy5 channel of the image, and the “measure” functionality was used to quantify the mean fluorescent intensity of the Cy5 within the bounds of the thresholded region.

### CNS and choroid plexus scRNA-seq sample preparation

Mice were perfused with heparinized (10 U/ml) DPBS and a single cell suspension of the entire brain (including ChPs) was generated using the adult brain dissociation kit as described earlier. The cells were then washed twice with cell-suspension buffer (FB0002440) to remove debris and counted on a hemocytometer. Cells were diluted to load 40,000 cells for encapsulation in a PIPseq T20 kit, as recommended by manufacturer with the intent of recovering 20,000 cells. Single cell libraries were then prepared using the v4.0 PIPseq T20 kits following manufacturer specifications. Sequencing was performed on an Illumina NovaSeq6000 PE150 at a depth of 400 million total reads, yielding a final depth of 20K reads per cell.

In an independent experiment under the same dosing conditions, choroid plexus tissue was isolated for scRNA-seq. We adapted a previously described method to generate a single cell suspension of ChP cells [37]. In brief, all ChPs (both laterals, 4^th^, and 3^rd^ ventricle ChPs) were pooled from N=4 mice injected with either L2-siRNA^PPIB^ or L2-siRNA^NTC^ and thoroughly minced with scissors in cold RPMI media. The tissue was further dissociated in digestion media for 30 minutes at 37 ^◦^C in an inverting tube rotator, with trituration every 10 minutes. The digestion media was composed of collagenase/dispase (1 mg/mL, Sigma 63837023) with DNAse (250 U/mL, Sigma 04536282001). Cells were pelleted post-digestion and washed in cell-suspension buffer. After passing cells through a 35 μm filter, cells were washed again and counted. The cells were then loaded into PIPseq T20 kits as described above.

### scRNA-seq data processing

Reads were processed using the PIPseeker alignment algorithm (v02.01.04), and the resulting count matrices were analyzed with standard methods in Seurat (v4). The datasets were merged without applying batch correction, and cells were filtered based on the number of features, retaining cells with 800–10,000 uniquely expressed genes. The counts were normalized, and the dataset was reduced to the top 2,000 variable genes. Data scaling was performed, followed by dimensionality reduction (PCA and UMAP) based on the first 50 principal components to facilitate data visualization. Cells were clustered using the Louvain algorithm and annotated according to standard marker genes from the mouse brain atlas [38]. Average expression levels of Ppib were computed using Seurat’s AverageExpression() function and reported for each cluster in biological replicates. Wilcoxon statistical test was used to evaluate *Ppib* as the differentially expressed gene within each cell type. Bonferroni’s correction for multiple comparison’s was applied to account for the number of genes being computed for differential expression.

## RESULTS

### L2-siRNA accumulates in brain endothelial cells after intravenous administration

To assess delivery to brain endothelial cells, we compared biodistribution and cell-specific uptake of L2-siRNA and an unconjugated siRNA lacking lipidation. Both the L2-siRNA and control siRNA structures were synthesized with modified chemistries at the 2L ribose position (alternating 2LFluoro and 2LOMe in a zipper pattern) and in the backbone (the two terminal bases are linked with phosphorothioate bonds at both ends of both strands of the duplex), while the 5L antisense strand contains a vinyl phosphonate. As previously described [32–34], when siRNA is conjugated with the lipids to form L2-siRNA, the divalent 18-carbon lipid tails are spaced from the branch point by ethylene glycol blocks, each linked through phosphorothioate bonds for a total of 18 ethylene glycols (Fig. 1A). We injected 20 mg/kg of Cy5-labeled oligonucleotide and assessed delivery with flow cytometry to quantify cell-specific uptake and with histology to visualize CNS distribution (Fig. 1B). Endothelial cells were gated as positive for CD31 and negative for O1, which was used as a counterstain to exclude oligodendrocytes (Supplemental Fig. 1) [39]. Within this CD31^pos^/O1^neg^ population, L2-siRNA exhibited potent uptake, as measured by Cy5 intensity (Fig. 1C). While L2-siRNA achieved greater delivery than siRNA in all cell types examined, the accumulation in CD31^pos^ brain endothelial cells is most striking, where L2-siRNA is detected in ∼100% of these cells, in contrast to the unconjugated siRNA, which was only present in ∼25% of this population (Fig. 1D).

**Figure 1:**
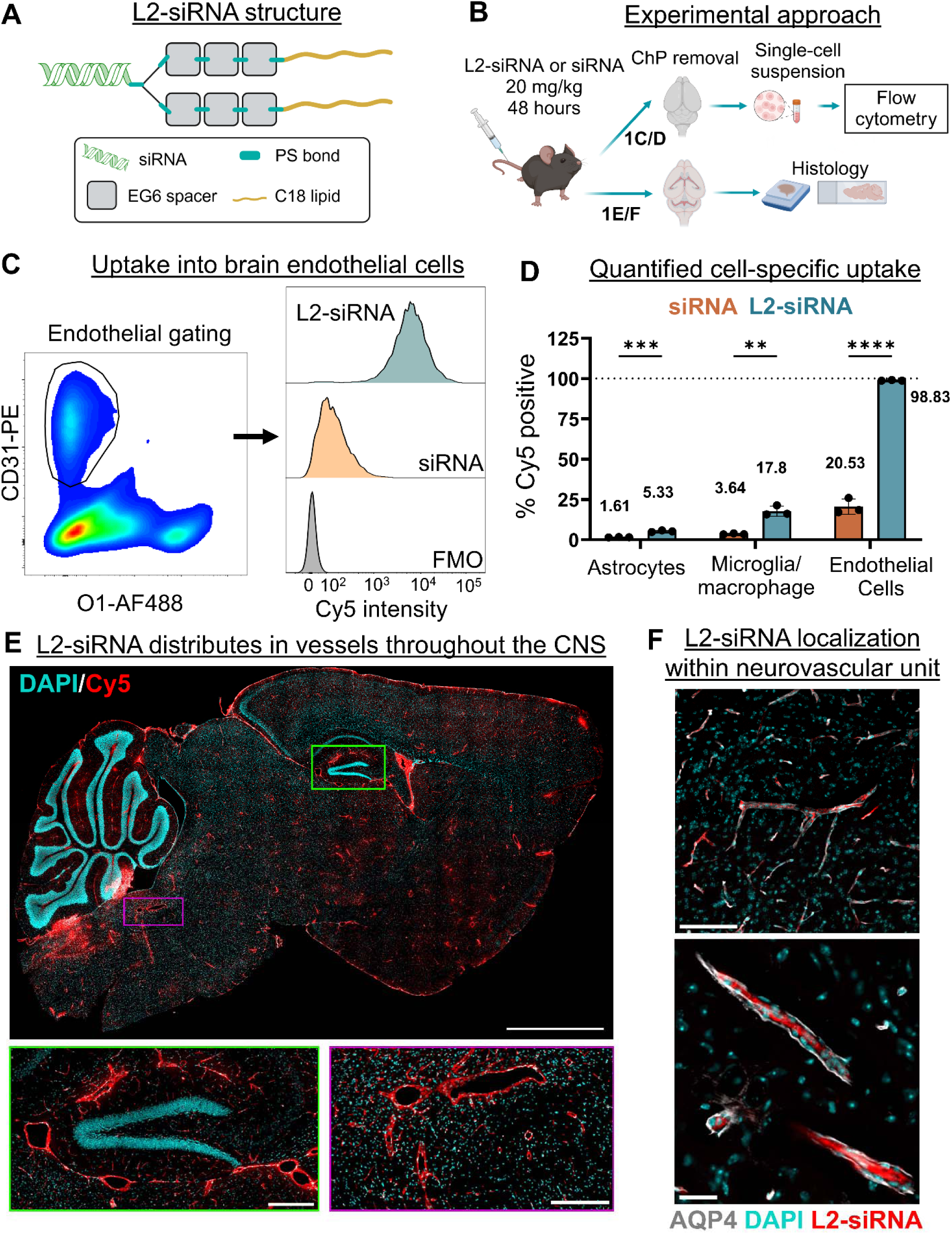
L2-siRNA accumulates in brain endothelial cells. A. Schematic of L2-siRNA structure with chemical modifications. B. Experimental design to assess endothelial uptake of L2-siRNA. Mice are injected with 20 mg/kg of Cy5 labeled L2-siRNA or siRNA, and after 48 hours, delivery is assessed by flow cytometry and histology. C. Endothelial cells are gated as CD31^Pos^/O1^Neg^, and Cy5 intensity is examined in this population compared to a fluorescence-minus-one (FMO) control. AF488 = Alexa Fluor 488, PE = phycoerythrin. D. Cell-specific antibodies are used to identify CNS populations of interest. Uptake of siRNA and L2-siRNA is quantified as percent positive based on FMO controls (i.e. brain cells from an uninjected mouse). Statistics are computed as multiple t-tests (Holm-Sidak correction) from N=3 mice, and data is presented as mean ± SD (* p<0.05, ** p<0.01, *** p<0.001, **** p<0.0001). Gating scheme and representative plots are depicted in Supplemental Fig. 1. Mean percent positive is shown above each respective bar. E. L2-siRNA distribution throughout the CNS (scale bar = 2 mm). Delivery to the brainstem (bottom right, scale bar = 250 µm) and hippocampus (bottom left, scale bar = 200 µm) highlight association with structures resembling blood vessels. Section thickness = 20 µm. F. L2-siRNA delivery is confined to the area enclosed by AQP4+ astrocyte endfeet. Representative thalamic vessels shown at low (upper panel, scale bar = 100 µm) and high magnification (bottom, scale bar = 25 µm).

L2-siRNA delivery to the vascular endothelium was further corroborated using a histological approach. After 48 hours, L2-siRNA is retained in a vessel-like pattern throughout the entire parenchyma. Enlarged images of the brainstem and hippocampus highlight distribution into both large and small vessels (Fig. 1E). A saline-injected negative control demonstrates minimal background signal, further supporting the observed vessel-association with L2-siRNA (Supplemental Fig. 2A). Similar distribution is observed in the blood-spinal cord barrier, a morphological correlate of the BBB that possesses analogous barrier properties (Supplemental Fig. 3). To bolster these findings, we more rigorously characterized the localization of L2-siRNA within the neurovascular unit - a structure comprised of an endothelial cell, endothelial basement membrane, pericyte, astrocytic/pericyte basement membrane, and the outermost glia limitans layer [40]. Staining for the glia limitans with AQP4 revealed that L2-siRNA is contained within this boundary and has not penetrated the parenchyma (Fig. 1F). Due to washout of lipophilic conjugates when permeabilizing the membrane with detergents, we limited our immunofluorescent staining to surface markers, such as AQP4, that do not require permeabilization. Taken all together these data collectively highlight widespread yet selective accumulation of L2-siRNA into brain endothelial cells after intravenous delivery.

### L2-siRNA accumulates in the choroid plexus after intravenous administration

Given that systemically delivered compounds should also reach the ChP through fenestrated capillaries, we assessed L2-siRNA distribution in ChPs relative to siRNA. Cy5-tagged compounds were intravenously injected at 20 mg/kg, and the ChPs were extracted after 48 hours for histological analyses. Compared to siRNA, L2-siRNA achieved greater absolute delivery both qualitatively (representative image shown in Fig. 2A) and quantitatively throughout all ChPs (Fig. 2B). The quantitative analysis is described in Supplemental Fig. 4. L2-siRNA was effectively delivered to each of the choroid plexuses (i.e. lateral, 4^th^, 3^rd^).

**Figure 2:**
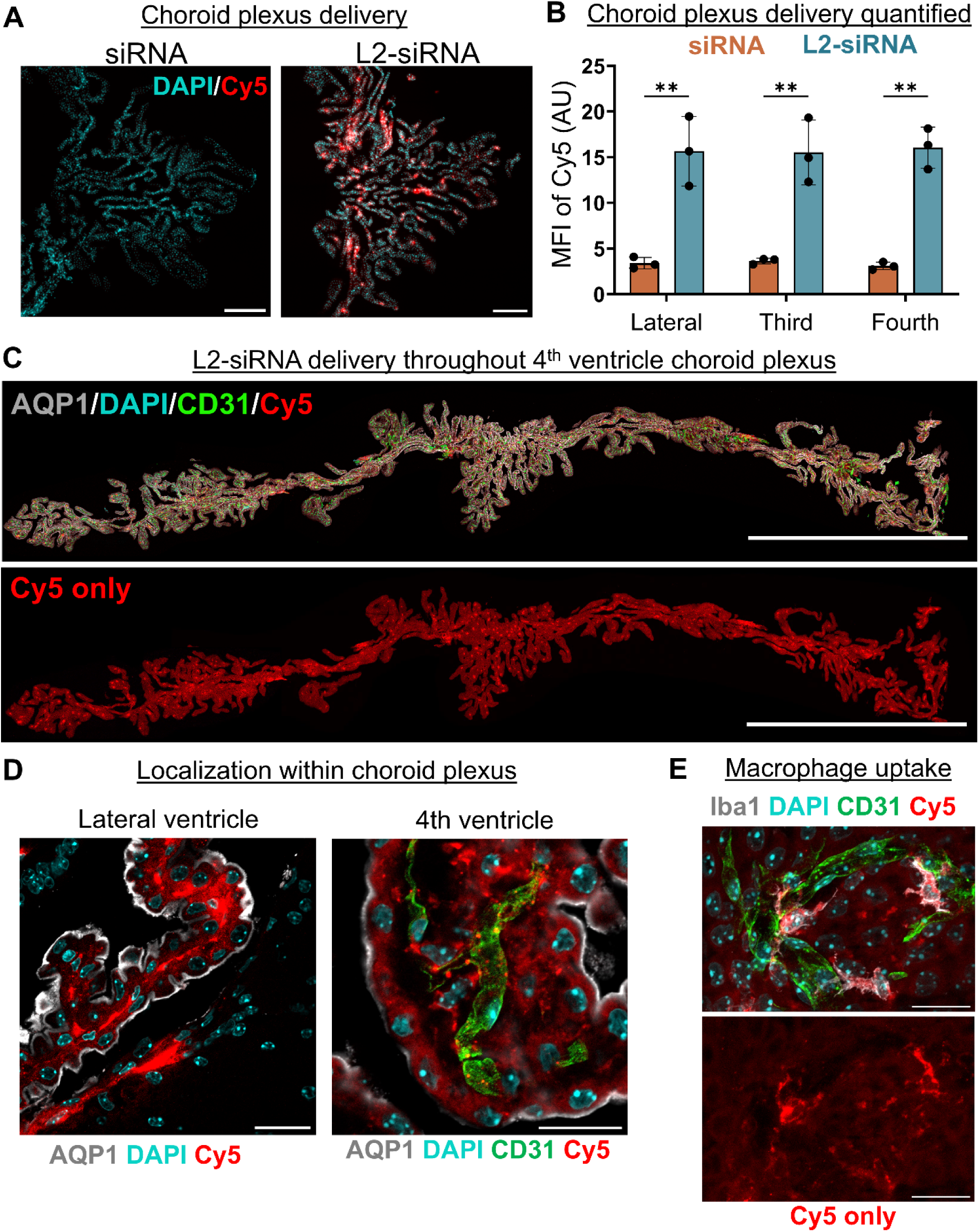
L2-siRNA accumulates in choroid plexus within diverse cell populations. A. Representative images of delivery to the 4^th^ ventricle choroid plexus 48 hours after a 20 mg/kg intravenous injection of Cy5-labeled siRNA and L2-siRNA. Scale bar = 200 µm. B. Quantification of Cy5 mean fluorescence intensity (MFI) in each choroid plexus. Data are presented as mean ± SD from N=3 mice. Statistics computed as multiple unpaired two-tailed t-tests (Holm-Sidak correction; ** p<0.01, *** p<0.001, **** p<0.0001). Quantitation approach is outlined in Supplemental Fig. 3. C. The 4^th^ ventricle ChP is whole mounted (confocal z-stack projection, scale bar = 2 mm) and labeled with aquaporin 1 (AQP1) staining the apical side of the epithelium and CD31 staining the vasculature. D. L2-siRNA spatial distribution in choroid plexus is shown at the base of the lateral ventricle (left, scale bar = 25 µm) and in a 4^th^ ventricle explanted choroid plexus (right, scale bar = 20 µm). E. Fluorescent L2-siRNA signal colocalizes with Iba1+ macrophages. Scale bar = 20 µm.

Since L2-siRNA delivery was not homogenously dispersed *within* each ChP, we next investigated L2-siRNA localization to different regions and cell types. Delivery was present throughout the entire structure, as demonstrated in a whole mount image of the 4^th^ ventricle ChP (Fig. 2C). We employed cell-specific immunostaining to more clearly identify regions and cell types with preferential L2-siRNA accumulation. CPECs are the most abundant population in the ChP and perform essential functions in regulating the composition of CSF. Staining the apical (CSF-facing) membrane of CPECs with AQP1 highlights widespread uptake into these barrier-forming cells (Fig. 2D). L2-siRNA can also be visualized in the stromal space, which lies between the endothelium (CD31+) and epithelium (AQP1+) (Fig. 2D). Lastly, to investigate which cell types had greatest uptake (observed as bright punctate signal in the explant image), we stained ChP macrophages with an anti-Iba1 antibody. We found that these macrophages exhibited extensive uptake as evidenced by colocalization between Iba1+ macrophages and Cy5 within the stromal space where these cells reside (Fig. 2E). Overall, we highlight transport to all choroid plexuses and cell types examined, including macrophages and CPECs.

### L2-siRNA promotes sustained gene silencing in brain barrier cells

Based on the promising delivery of L2-siRNA to brain barrier cells, we next characterized the potency and temporal kinetics of target gene silencing. We chose *Ppib* as a proof-of-concept gene target that is ubiquitously expressed by CNS cells (including brain barrier populations) [41] and for which a well-validated siRNA sequence had been identified [42]. Knockdown was assessed in ChPs by manually dissecting these regions and performing RT-qPCR analysis. To isolate brain endothelial cells, the brains (excluding ChPs) were dissociated into single cells and sorted for CD31^pos^ endothelial cells. Using RT-qPCR, we validated that these isolated cells were enriched for expression of the endothelial cell marker *Cldn5* and lacked contamination from astrocytes expressing *Gfap* and microglia expressing *Iba1* (Supplemental Fig. 5). A non-targeting control (NTC) siRNA (termed L2-siRNA^NTC^) was used as the primary negative control for all gene silencing studies and did not change *Ppib* expression in brain endothelial cells compared to an uninjected control (Supplemental Fig. 6).

We assessed knockdown at three timepoints (8, 15, 30 days) following a single 20 mg/kg injection (Fig. 3A). Interestingly, L2-siRNA targeting *Ppib* (L2-siRNA^PPIB^) potentiated robust gene silencing in brain endothelial cells at 8 and 15 days (∼50%) but returns to basal levels by 30 days (Fig. 3B). In contrast, CD31^neg^ cells, comprising all isolated CNS cells except endothelial and choroid plexus cells, did not exhibit detectable knockdown, suggesting that gene inhibition is more effective in the endothelium than parenchyma (Fig. 3C). Bulk choroid plexus knockdown kinetics were different than the brain endothelium. Intriguingly, the lateral ventricle ChP achieved peak knockdown 30 days after injection, highlighting a prolonged duration of gene silencing from a single intravenous injection (Fig. 3D). Similarly, the fourth ventricle ChP achieves moderate knockdown by 8 days, which persisted to 30 days, the longest time examined (Fig. 3E). Collectively, these data showcase L2-siRNA as a platform for mediating gene inhibition in both the BBB and BCSFB.

**Figure 3:**
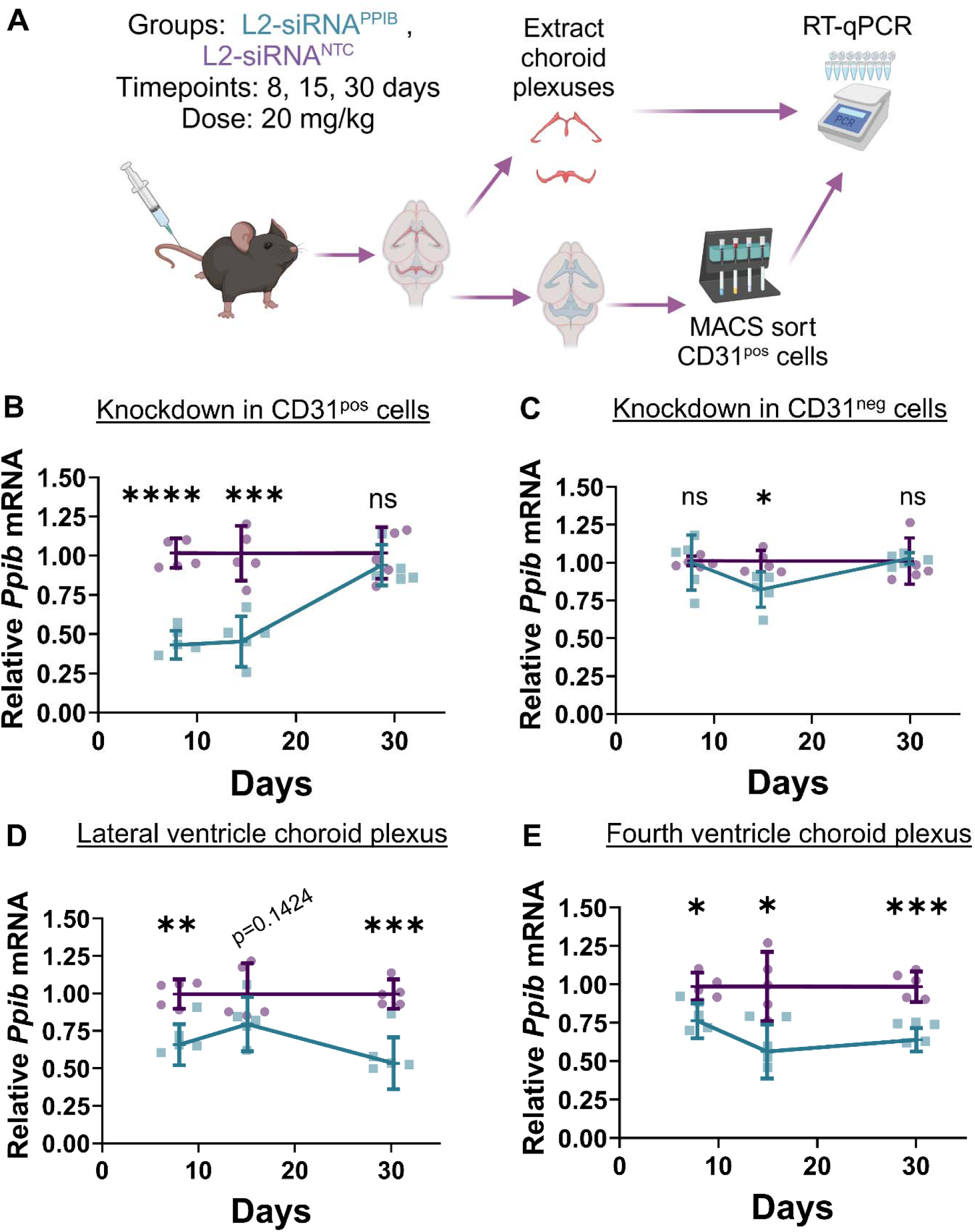
L2-siRNA promotes sustained gene silencing in brain barrier cells and tissues. A. Experimental design to assess *Ppib* silencing in brain endothelial cells and whole choroid plexuses. Knockdown was assessed at three timepoints (8, 15, 30 days) after a single 20 mg/kg intravenous injection. B. Time course of mRNA knockdown in CD31^pos^ endothelial cells as assessed by RT-qPCR. C. Time course of mRNA knockdown in non-endothelial CNS cells (CD31^neg^). D. Gene silencing in the choroid plexuses of the two lateral ventricles combined. E. Gene silencing in the choroid plexus of the fourth ventricle. For all panels, N=5 biological replicates (i.e. individual mice) were used for each timepoint, and data are presented as mean ± SD. Unpaired two-tailed t-tests for each timepoint normalized to L2-siRNANTC (* p<0.05, ** p<0.01, *** p<0.001, **** p<0.0001, ns – not significant).

### Cell-specific knockdown across the endothelium and choroid plexus

We next sought to understand knockdown in different vessel types, whether parenchymal cells experience gene silencing, and the cell type-dependent silencing activity within the ChP. To that end, we employed single cell RNA sequencing (scRNA-seq) to identify specific cell populations and then assessed *Ppib* knockdown within these cell types similar to our prior work [34]. The analysis was performed on dissociated brains 15 days after intravenous injection of L2-siRNA^PPIB^ compared to L2-siRNA^NTC^ (Fig. 4A). To confirm *Ppib* knockdown in this experiment, the CD31^Pos^ cells not used for scRNA-seq were sorted for RT-qPCR analysis. Consistent with prior knockdown studies, L2-siRNA^PPIB^ showed 50% knockdown in CD31^Pos^ cells without knocking down *Ppib* in the CD31^Neg^ fraction (Supplemental Fig. 7). In the scRNA-seq samples, endothelial cells were segmented across the arterial – venous tree based on established markers, including *Bmx, Sema3g* (Arterial), *Slc16a2, Mfsd2a* (Capillary and Venous), *Slc3a85* (Venous), *Vwf* (Venous and Arterial) (Fig. 4B, Supplemental Fig. 8) [41]. Amongst these vascular cells, statistically significant *Ppib* knockdown was observed exclusively in capillary and venous endothelial cells, with venous endothelial cells exhibiting the greatest gene inhibition (Fig. 4C). No parenchymal cells exhibited statistically significant gene silencing (Fig. 4C). The only other cell population differentially expressing *Ppib* was CPECs, consistent with the selective delivery of L2-siRNA to brain barriers. The reduction in *Ppib* is also visualized as a bulk shift in the large peak of the ridgeline plots (Fig. 4D). These plots were helpful as they reveal a bimodal distribution, which is expected for any gene because sequencing coverage results in a Poisson distribution with a high proportion of zeroes [43,44].

**Figure 4:**
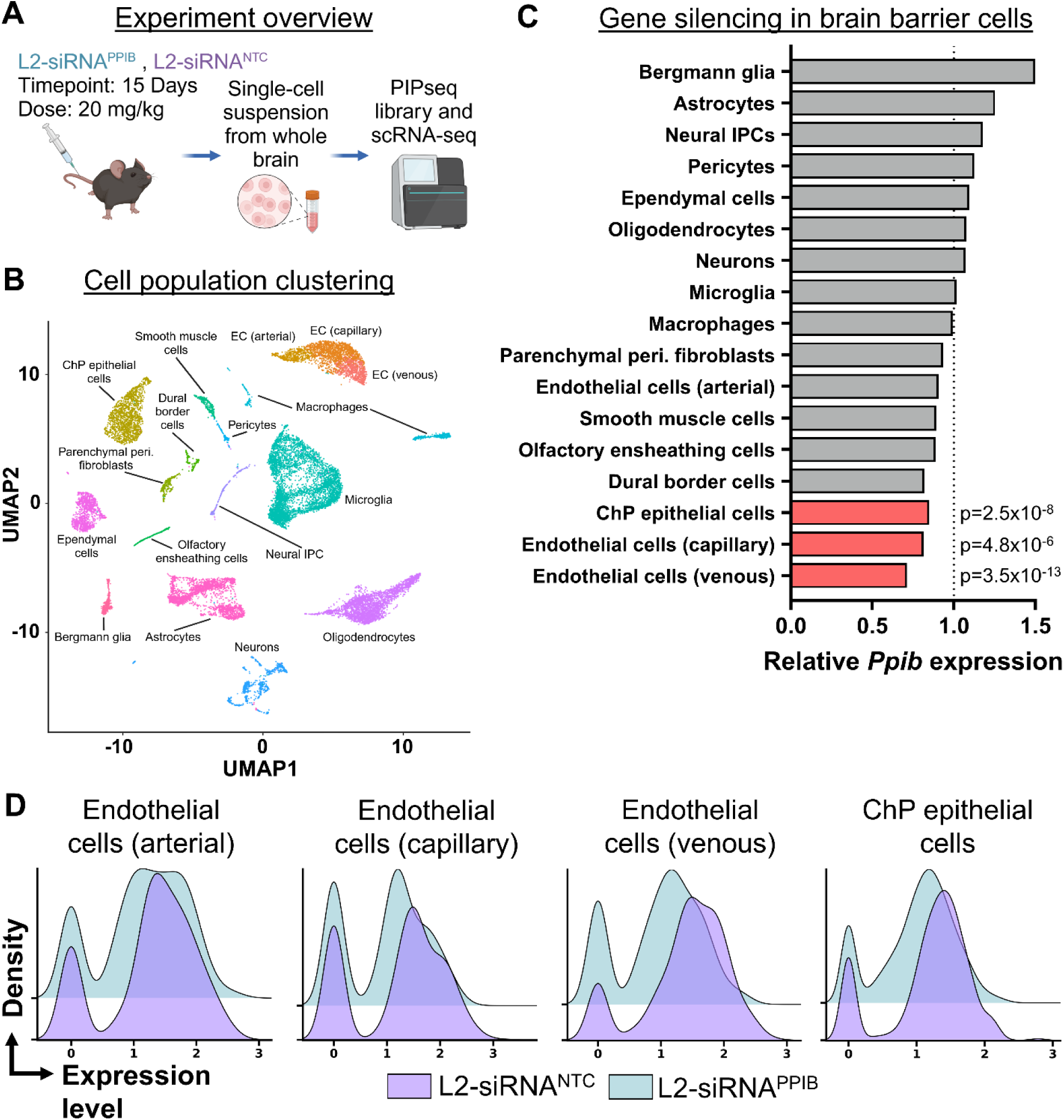
scRNA-seq analysis of gene inhibition across whole brain. A. Experimental design to assess cell-specific silencing. Mice were intravenously injected with 20 mg/kg L2-siRNA^PPIB^ or L2-siRNA^NTC^. After 15 days, each brain was dissociated into single cells and sequenced. B. Dimension reduction plot showing clustering and annotation of identified cell populations. C. *Ppib* expression for each cell population in the L2-siRNA^PPIB^ samples relative to the L2-siRNA^NTC^ samples. Cell types with statistically significant reduction in *Ppib* are depicted in red (Wilcoxon test with Bonferroni’s correction for multiple comparisons). D. Ridgeline plots showing distribution of *Ppib* expression in brain barrier cells.

To more closely examine cell-type specific knockdown at the BCSFB, we performed an additional scRNAseq experiment on pooled ChP samples. Consistent with whole brain sequencing, we detected target gene knockdown in CPECs (Supplemental Fig. 9). We also identified endothelial, immune, mesenchymal, and glial populations; however, due to the relative abundance of epithelial cells, the number of cells captured in these other cell types was underpowered to compute significance.

### Peripheral biodistribution and gene silencing efficacy of L2-siRNA

Beyond brain barriers, intravenous delivery of L2-siRNA provides an opportunity to examine transport to anatomical sites outside the CNS. Further, characterizing knockdown activity in peripheral organs provides a framework for considering off-target effects of L2-siRNA when selecting downstream therapeutic targets relevant to brain barriers. As part of a detailed examination, we assessed organ biodistribution at 48 hours as well as delivery and gene silencing kinetics over a 30-day course matching the one used in Figure 3 with the same 20 mg/kg single injection.

We analyzed five major organs relevant to siRNA transport – liver, heart, spleen, lung, and kidney. The highest L2-siRNA accumulation was observed 8 days post-injection in the liver, as evidenced by histology and a peptide nucleic acid (PNA) probe-based assay for siRNA detection. Accordingly, the liver achieved the greatest gene silencing out of all the organs; ∼70% decrease in *Ppib* at 30 days (Fig. 5A). L2-siRNA was also transported to the heart and lungs, exhibiting non-homogenous distribution in these tissues and a gradual clearance over time (Fig. 5B,C). Correspondingly, knockdown in the lungs and heart was greatest at 8 days and then gradually lessened. In contrast, the spleen experienced considerable accumulation, particularly in the red pulp, but this uptake did not correspond with productive gene inhibition (Fig. 4D). Consistent with prior studies, albumin-binding L2-siRNA does not accumulate or yield any gene inhibition in the kidney (Fig. 4E), suggesting that the lipid conjugate may remain in circulation by escaping renal clearance pathways.

**Figure 5:**
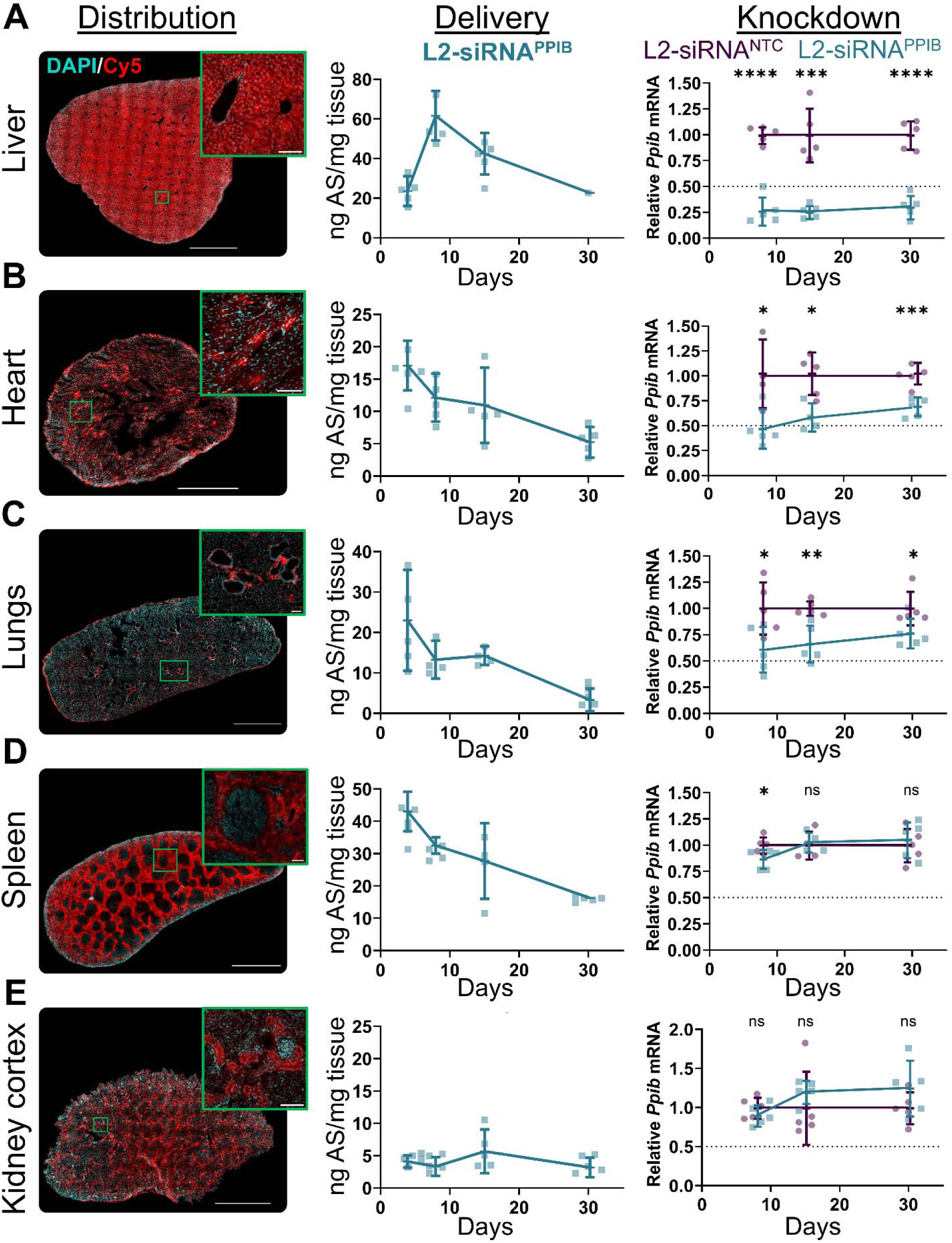
L2-siRNA distribution and gene silencing in peripheral organs. Biodistribution, siRNA accumulation, and *Ppib* knockdown measured in each organ after a 20 mg/kg intravenous injection of L2-siRNA. Histology was performed 48 hours after injection. Delivery and knockdown were assessed at the described time points. N=5 mice in each condition, represented as mean ± SD. All panels are annotated with unpaired two-tailed t-tests for each timepoint normalized to L2-siRNANTC (* p<0.05, ** p<0.01, *** p<0.001, **** p<0.0001, ns – not significant). The representative images of each organ were imaged identically, however, the brightness of the liver and spleen was reduced in processing to prevent saturation. Scale bar = 100 μm for all insets and scale bar = 2 mm for all organs except the heart, which is 1 mm. A. Distribution and gene silencing in the median lobe of the mouse liver. For histology, the median lobe was used in its entirety. For gene silencing, hepatic tissue was biopsy punched between the center and edge. B. Distribution and gene silencing in the left ventricle of the heart. A transverse section of the ventricle is shown for histology. Samples for delivery and gene silencing were collected from the outer wall of the left ventricle. C. Distribution and gene silencing in the lung. Samples were taken from the left lung. D. Distribution and gene silencing in the spleen. Delivery and gene silencing were assessed in samples containing both white and red pulp. E. Distribution and gene silencing in the kidney. Histology was assessed in kidney cortex. Delivery and gene silencing were assessed in samples containing both the cortex and medulla.

## DISCUSSION

Dysfunctional brain barriers contribute to the pathophysiology of chronic CNS diseases, yet few non-viral technologies are established that therapeutically target these interfaces. In this work, we established that a lipid-siRNA conjugate structurally optimized for albumin binding has great potential to be applied for therapeutic strategies that modulate gene expression in brain barriers. We highlight >50% target gene silencing in brain endothelial cells that is sustained for up to two weeks, and prolonged knockdown in bulk choroid plexus over a one-month period. Analysis of cell-specific knockdown further supported potent activity in barrier-forming cells (venous and capillary endothelium and CPECs) without affecting parenchymal cells. Here, we provide additional commentary on the mechanisms of delivery to blood-CNS barriers, kinetics of gene silencing, and activity along the arterio-venous spectrum.

Tissue distribution of lipid-siRNA conjugates is influenced by association with serum proteins and other components, and we hypothesized that reversible binding of L2-siRNA to albumin may facilitate brain vessel accumulation. Initial evidence for how protein binding can govern transport comes from first generation lipid-siRNA conjugates, such as cholesterol-siRNA, which bind to high-density lipoprotein (HDL) and are shuttled to the liver [45]. In terms of brain delivery, further investigation revealed that cholesterol-siRNA could achieve gene silencing in brain endothelial cells only when pre-complexed with HDL (not unbound, or with LDL), and parenchymal gene silencing was not assessed [46]. In contrast, pre-complexing cholesterol-siRNA with albumin steered delivery to extrahepatic sites such as the spleen and heart [45]. GLP-1 agonist Semaglutide (Ozempic) also leverages albumin binding to endow a long pharmacokinetic profile optimized for extracellular engagement of GLP-1R [47]. In the brain, delivery was restricted to circumventricular organs (brain regions lacking a BBB) where Semaglutide acted on neurons to promote satiation. Interestingly, although both L2-siRNA and Semaglutide bind albumin in the bloodstream, Semaglutide did not accumulate in brain endothelial cells [48]. This discrepancy may be governed by structural differences, as L2-siRNA albumin-binding and cell uptake properties are based more on hydrophobic interactions (all hydrocarbon lipid without ionizable acid groups) [32], suggesting that the L2-siRNA can release from albumin and interact with nearby endothelial cell membranes. Further investigation is warranted to understand if tuning the albumin affinity could be an effective strategy to promote brain barrier delivery more broadly.

One advantage of siRNA technology is the potential to generate sustained gene inhibition. The longevity of silencing depends on tissue type, cell composition, and injection route. In previous work, we demonstrated up to 5 months of bulk tissue CNS gene silencing for L2-siRNA injected directly into CSF [34]. The prolonged silencing is commonly attributed to the “endolysosomal depot” effect, which suggests that chemically stabilized siRNA is viable in acidic compartments, gradually leaking out into the cytoplasm and mediating RISC-induced mRNA degradation [49]. In this work, knockdown in brain endothelial cells was observed for 15 days but returned to baseline by one month. This time course is consistent with 7C1 endothelial-targeting nanoparticles in lung endothelial cells, which achieved rapid siRNA-mediated ICAM-2 knockdown but returned to basal expression levels by 28 days; notably this formulation did not exhibit gene silencing in brain endothelial cells [50,51]. In comparison to endothelial cells, we observed a greater duration of L2-siRNA silencing in choroid plexus (lateral and 4^th^). The difference in gene silencing kinetics is likely multi-factorial but may relate to cell turnover rates (CPEC vs. endothelial), active efflux transport within brain endothelial cells, or capacity for endosomal retention given that brain endothelial cells are very thin (∼200 nm). Also, as evidenced by Cy5 intensity on whole brain histology, the total delivery to the choroid plexus was far greater than to the endothelium. We posit that further modifications to L2-siRNA structure could be used to better understand these mechanisms and prolong gene silencing activity in future work.

Brain endothelial cells exhibit arteriovenous-specific functions relevant to transport and neurological disorders. For example, the post-capillary venule is the primary site for T-cell transmigration in multiple sclerosis (MS), driving white matter damage and debilitating symptoms [52]. Currently, FDA-approved therapies for MS deplete causal immune cells or block their interaction with the endothelium, leading to broad immunosuppression [53]. Targeting the venous endothelium with L2-siRNA to prevent immune cell extravasation into the CNS presents an opportunity for therapeutic intervention that may provide advantages over immunosuppression. Indeed, we observed the greatest targeting to the venous endothelium, which could be driven by slower blood flow hemodynamics increasing time for interactions between L2-siRNA and the endothelial membrane. Alternatively, the endothelial glycocalyx could influence delivery along the arterio-venous spectrum, as it repels negatively charged compounds and has greater coverage on arterioles than the capillary and venous endothelium [54]. The post-capillary venule has also been identified as the primary location for transcytosis of transferrin-receptor targeting nanoparticles, which achieved more transport across the post-capillary venule compared to capillaries and arterioles [55]. Thus, as a major anatomical site for transport into the CNS (molecular and cellular), the post-capillary venule is an excellent target for therapeutic intervention with L2-siRNA.

Collectively, we developed and characterized a new tool for manipulating gene expression in the barriers of the CNS. This work contributes to the emerging paradigm of therapeutically targeting the dysfunctional BBB or BCSFB.

## DATA AVAILABILITY

Raw sequencing data and processed Seurat objects are available at Array Express under accession number E-MTAB-14840. Full code has been made publicly available in a GitHub repository, which can be found at andrewkjar/Lipid-siRNA_systemic_scRNAseq.

## AUTHOR CONTRIBUTIONS

Conceptualization: AGS, KRS, CLD, ESL

Methodology: AGS and KRS performed the majority of experiments with assistance from NF (siRNA synthesis), AK (scRNA-seq), SL, JCP, AMA, DP, RPC, ENH, KAK, AW, ND

Supervision: ND, CLD, ESL Writing—original draft: AGS, KRS, CLD, ESL

Writing—review & editing: All authors reviewed and edited the manuscript

## ACKNOWLEDGEMENTS

Figures 1A,B; 3A; 4A,B; and Supplemental figures 7A,B,C and 8A,B were created with BioRender.com. Confocal microscopy was conducted in the Vanderbilt Cell Imaging Shared Resource core facility, which is supported in part by NIH grants P30 CA068485, P30 DK058404, and P30 EY008126. RNA sequencing was conducted in the Vanderbilt Technologies for Advanced Genomics core facility, which is supported in part by NIH grants 5UL1 RR024975, P30 CA068485, P30 EY008126, UL1 TR002243, and G20 RR030956. Flow cytometry experiments were performed in the VUMC Flow Cytometry Shared Resource, which is supported by the Vanderbilt Ingram Cancer Center (P30 CA68485) and the Vanderbilt Digestive Disease Research Center (DK058404). All LC-MS analyses were performed in the Mass Spectrometry Research Center at Vanderbilt University.

## FUNDING

This work was supported by the Chan Zuckerberg Initiative Ben Barres Early Career Acceleration Award grant [2019-191850 to ESL]; the National Institutes of Health [AG077807 to ESL & CLD, CA224241 to CLD]; Rita Allen Foundation Award [to ND]; NSF Graduate Research Fellowship Program [ENH & AK].

## COMPETING INTERESTS

ESL, CLD, and AGS are inventors on patent application number 18/737720 (applicant Vanderbilt University). This patent covers lipophilic siRNA conjugates for the treatment of CNS diseases.

