## Supplemental information for "Intravenous lipid-siRNA conjugate mediates gene silencing at the blood-brain barrier and blood-CSF barrier"

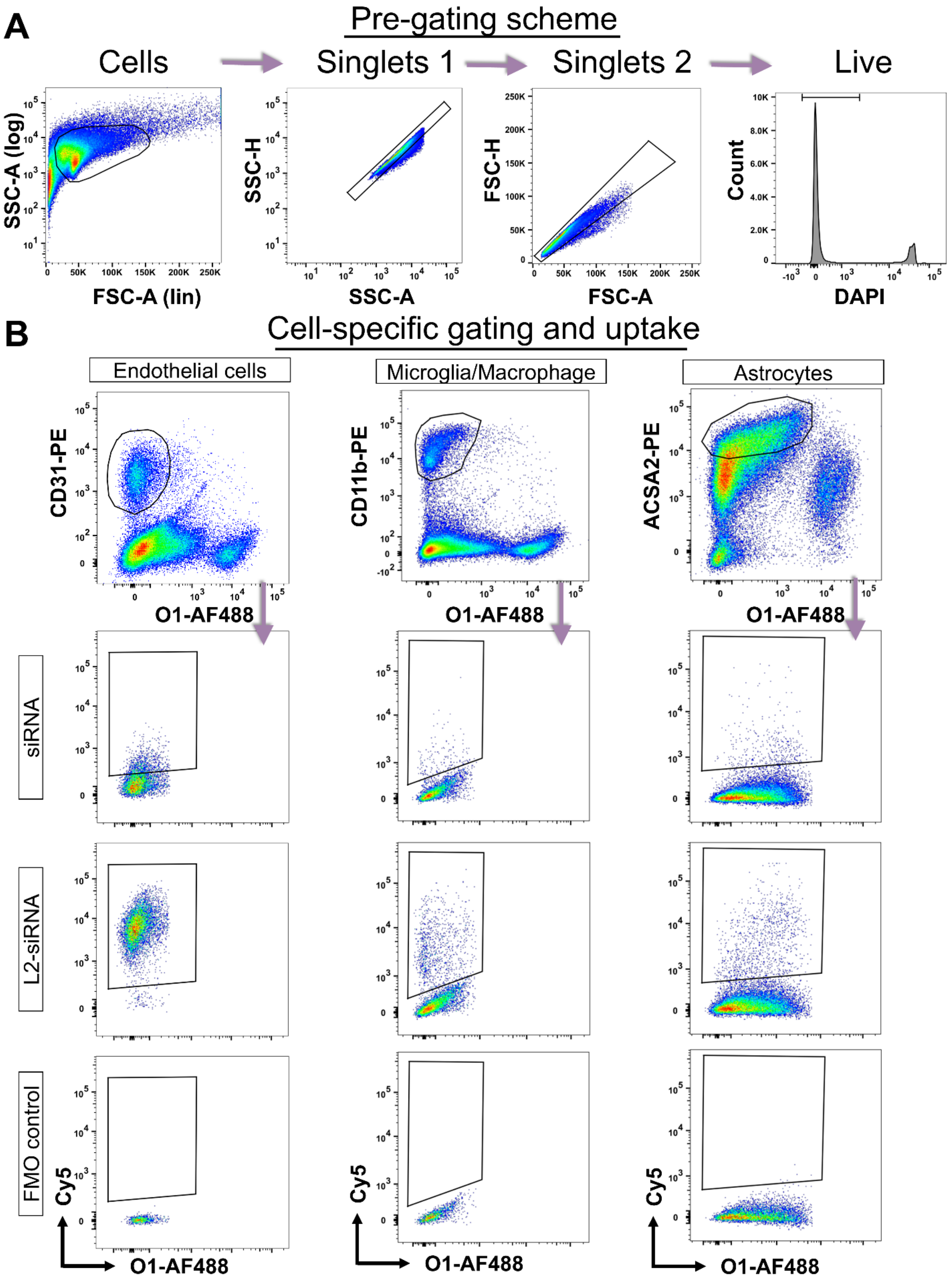

A
Pre-gating scheme
Cells
Singlets 1
Singlets 2
Live
SSC-A (log)
FSC-A (lin)
SSC-H
SSC-A
FSC-H
FSC-A
Count
DAPI
B
Cell-specific gating and uptake
Endothelial cells
Microglia/Macrophage
Astrocytes
CD31-PE
CD11b-PE
ACSA2-PE
O1-AF488
siRNA
L2-siRNA
FMO control
Cy5

**Supplemental Figure 1: Flow cytometry gating scheme for assessing cell-specific uptake**

A. Gating scheme applied to all samples to filter out debris, doublets, and dead cells.

B. Gating strategy for identifying endothelial cells, myeloid cells, and astrocytes based on validated markers. Cy5 positive events are gated using a fluorescence-minus one (FMO) control for each cell type. AF488 = Alexa Fluor 488, PE = phycoerythrin, FSC = forward scatter, SSC = side scatter.

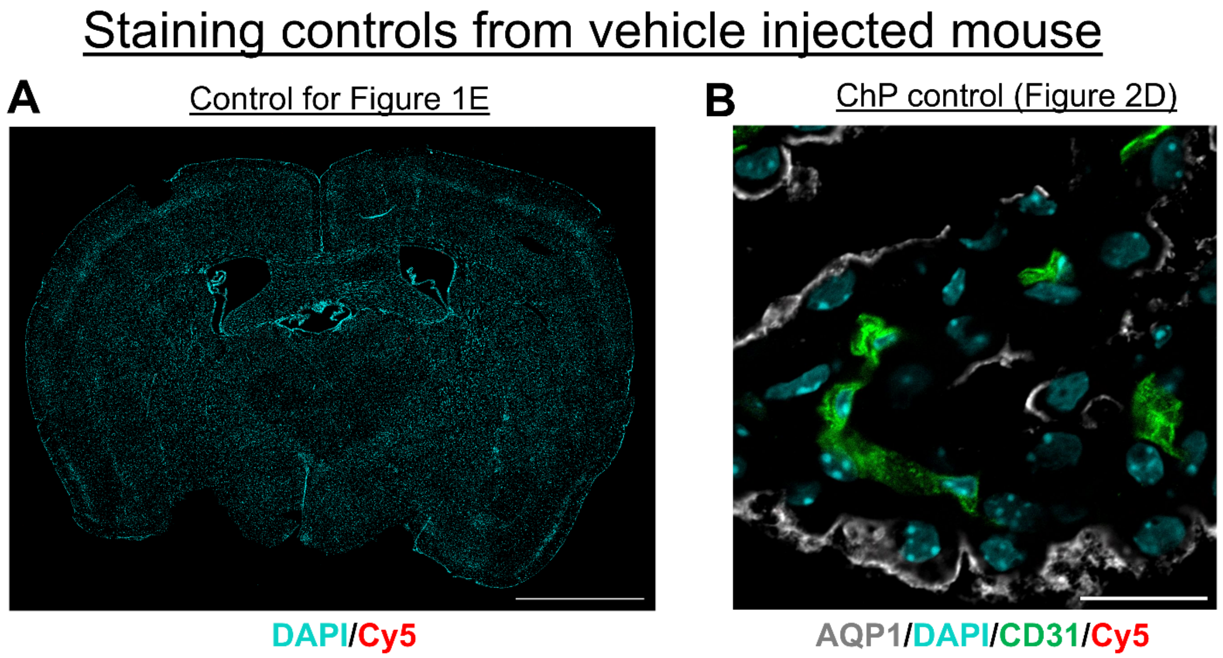


**Supplemental Figure 2: Staining controls to support biodistribution assessments**

Samples were prepared under the same conditions as experimental groups, except arising from a mouse injected with saline. (**A**) Coronal section processed and imaged identically to L2-siRNA sagittal section in Figure 1E. Scale bar = 2 mm. (**B**) Confocal image control is stained with the same antibodies as Figure 2D, and no Cy5 background signal was detected. Scale bar = 20 µm.

### Spinal cord biodistribution

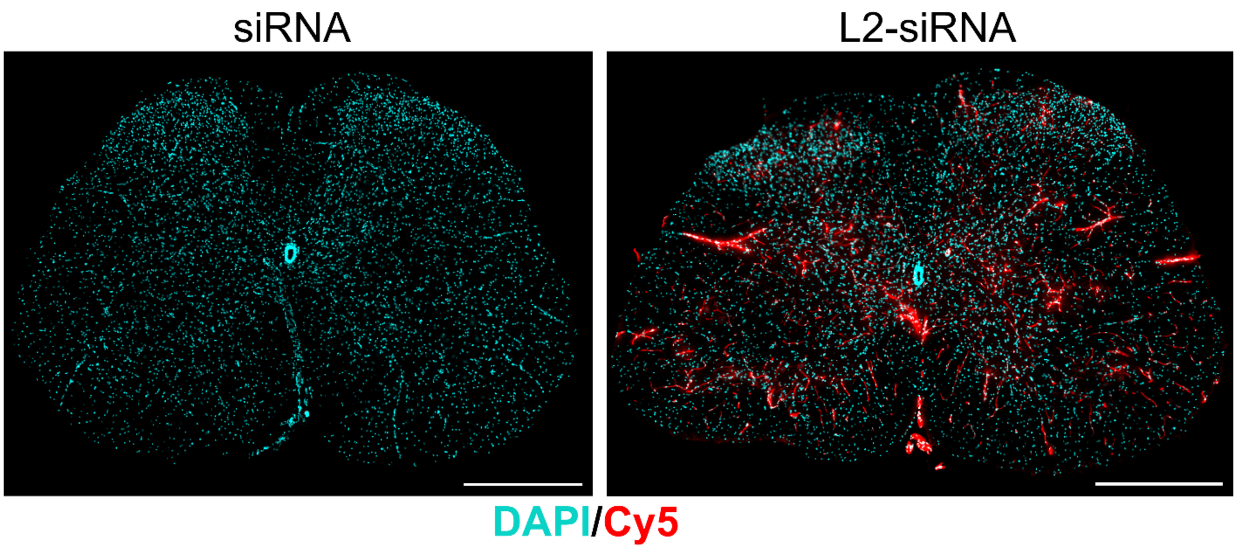


**Supplemental Figure 3: Spinal cord biodistribution**

Delivery to the spinal cord was assessed under the same conditions as brain biodistribution (20 mg/kg, 48 hours). 20 μm section thickness, scale bars = 2 mm.

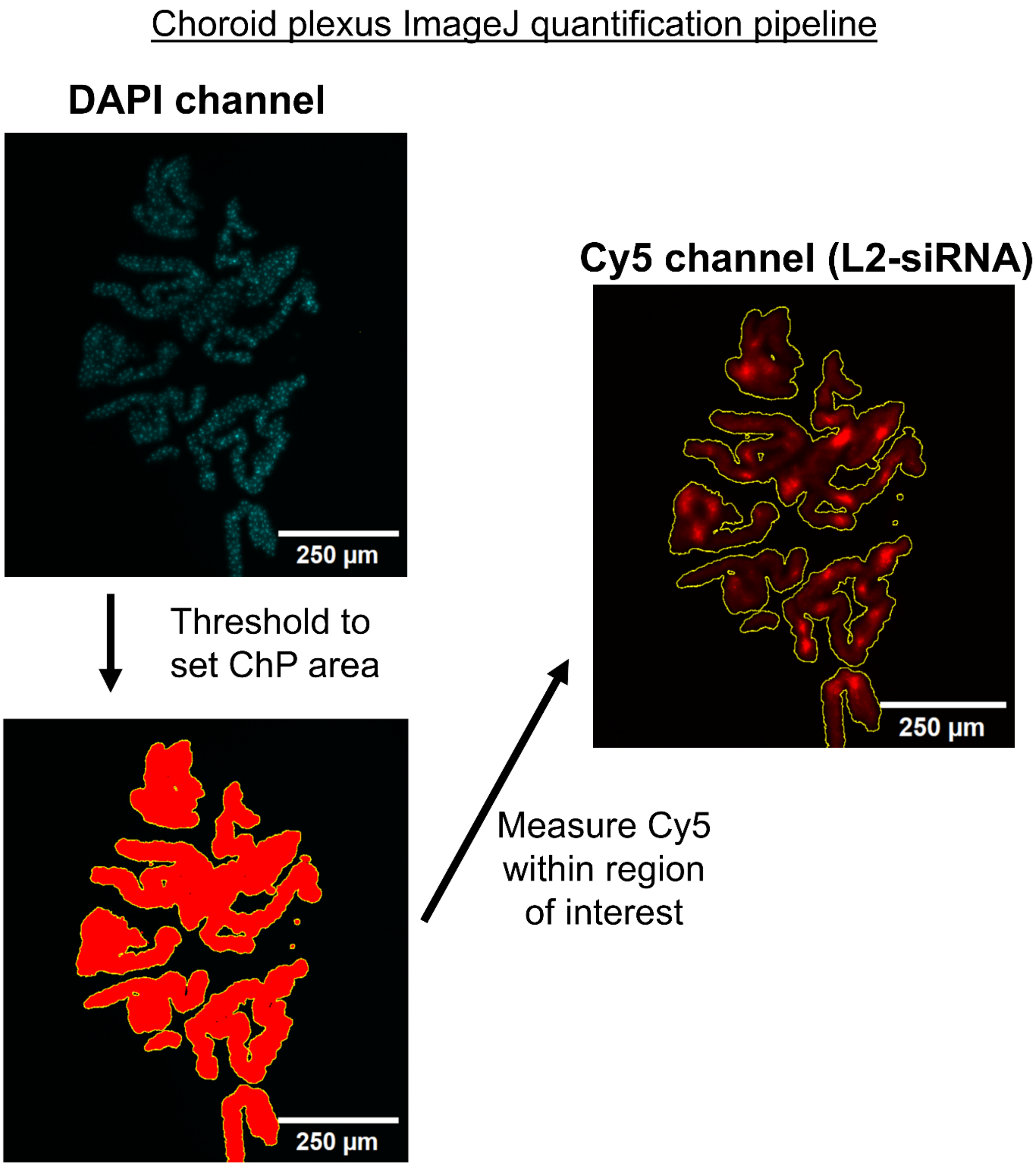


**Supplemental Figure 4: Image analysis pipeline for choroid plexus biodistribution.**

Representative image of a 20 µm section of the 4th ventricle choroid plexus displaying only the DAPI channel. Thresholding is applied on the DAPI channel with the Triangle algorithm in ImageJ. The thresholded area is then selected as an ROI and applied to the Cy5 channel to measure the mean fluorescent intensity of Cy5 in the bounded area.

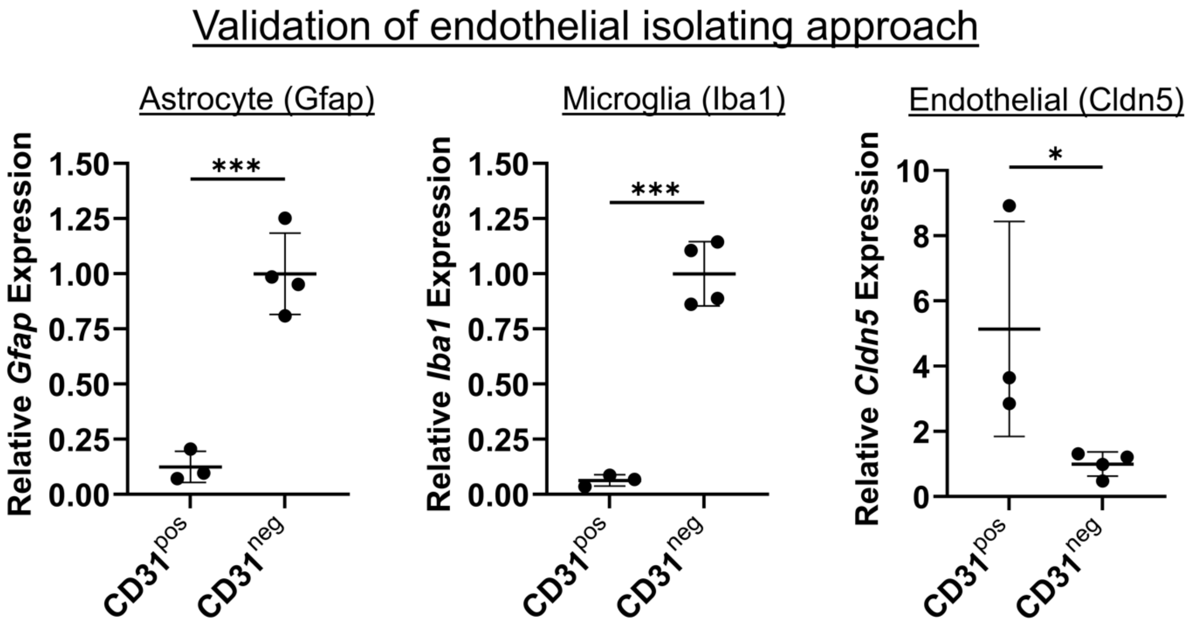


**Supplemental Figure 5: Validation of endothelial cell enrichment with anti-CD31 beads**

Brains from uninjected mice were sorted into CD31$^{pos}$ and CD31$^{neg}$ samples and analyzed with RT-qPCR. Canonical cell-specific markers show that the CD31$^{pos}$ cells are enriched for *Cldn5* expression and de-enriched for *Gfap* and *Iba1* expression. Data are normalized to CD31$^{neg}$ cells, and unpaired two-tailed t-tests were performed for each gene (N=3-4 replicates, where each data point is from cells isolated from a different mouse, mean ± SD shown; * $p<0.05$, *** $p<0.001$).

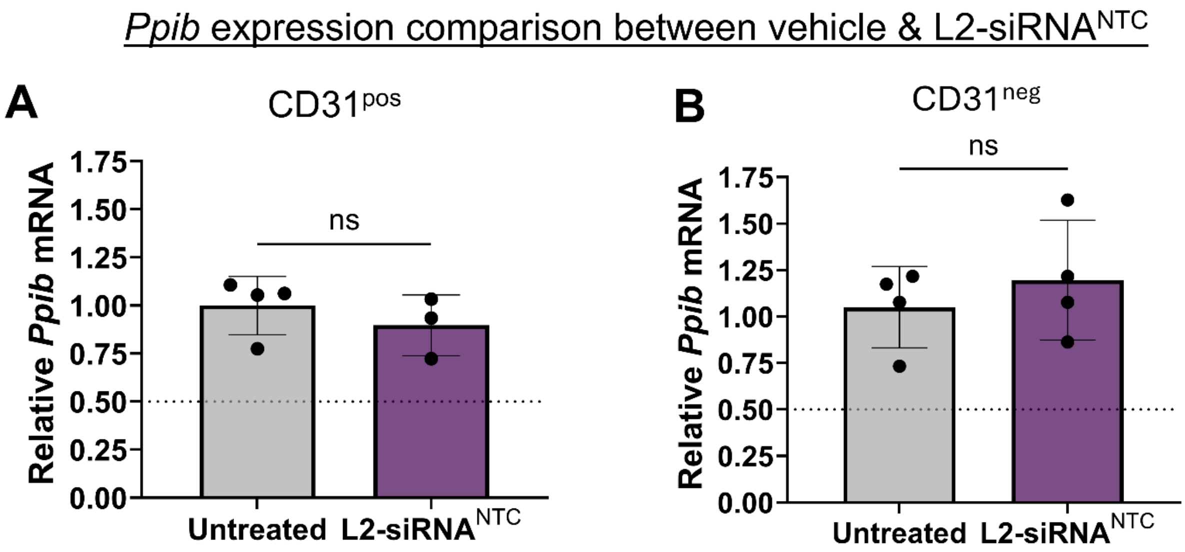


**Supplemental Figure 6: Evaluating *Ppib* expression in L2-siRNA$^{NTC}$-treated cells relative to uninjected mice**

CD31$^{pos}$ and CD31$^{neg}$ cells were analyzed for *Ppib* expression via RT-qPCR, either 48 hours after injection with 20 mg/kg L2-siRNA$^{NTC}$ or from an uninjected control. Each data point represents one mouse and is arbitrarily normalized to the uninjected cohort average. The mean and SD are displayed with the results of unpaired, two-tailed t-tests (ns – not significant).

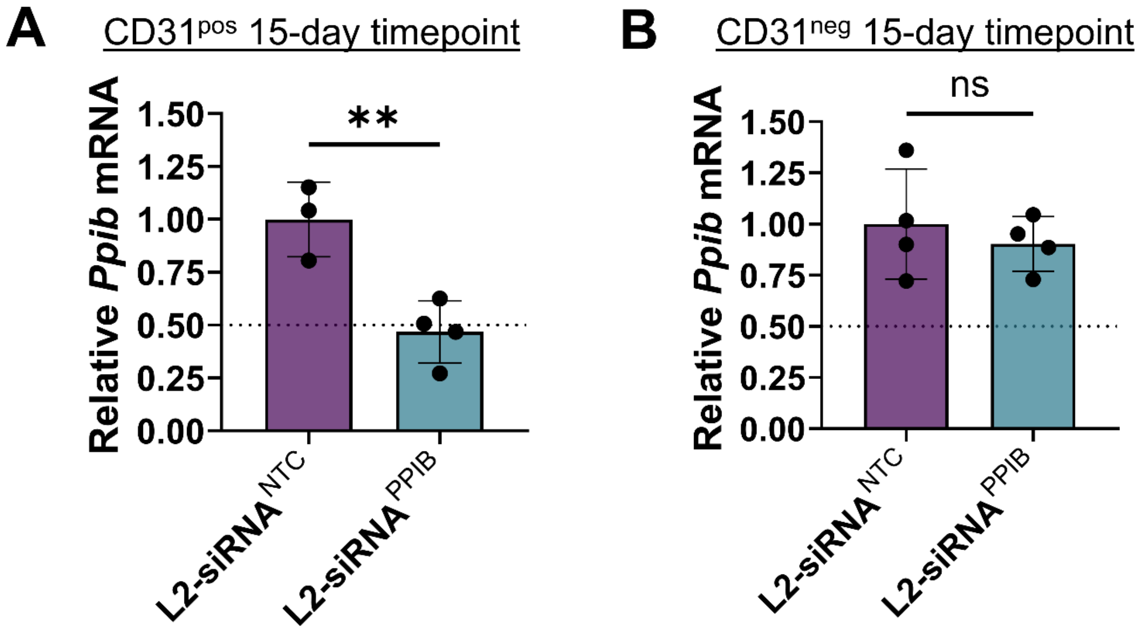


**Supplemental Figure 7: Additional independent experiment to validate *Ppib* inhibition in CD31$^{pos}$ brain endothelial cells in response to L2-siRNA$^{PPIB}$**

To demonstrate reproducibility, an independent study was performed to assess endothelial and parenchymal *Ppib* knockdown. Gene silencing was examined 15 days post 20 mg/kg IV injection of L2-siRNA $^{PPIB}$ or L2-siRNA $^{NTC}$. (**A**) Relative *Ppib* mRNA in CD31$^{pos}$ cells. (**B**) Relative *Ppib* mRNA in CD31$^{neg}$ cells. Each data point represents one mouse (N=3-4), and data are normalized to L2-siRNA$^{NTC}$. Significance was determined by unpaired two-tailed t-tests (** p<0.01, ns – not significant).

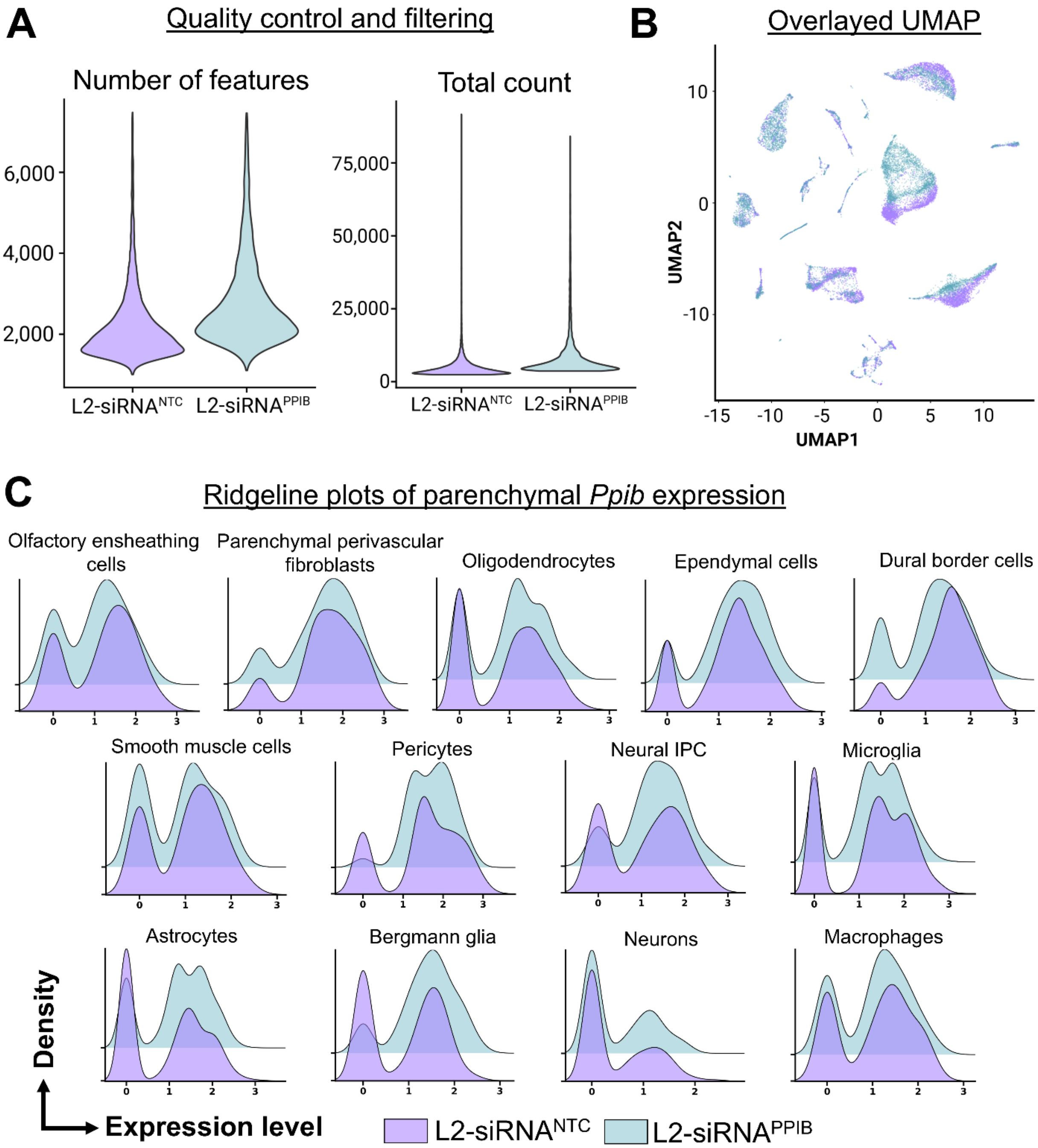
A
Quality control and filtering
Number of features
Total count
L2-siRNA^NTC
L2-siRNA^PPIB
B
Overlayed UMAP
UMAP1
UMAP2
C
Ridgeline plots of parenchymal *Ppib* expression
Olfactory ensheathing cells
Parenchymal perivascular fibroblasts
Oligodendrocytes
Ependymal cells
Dural border cells
Smooth muscle cells
Pericytes
Neural IPC
Microglia
Astrocytes
Bergmann glia
Neurons
Macrophages
Density
Expression level
L2-siRNA^NTC
L2-siRNA^PPIB

# D

#### Cell type annotation

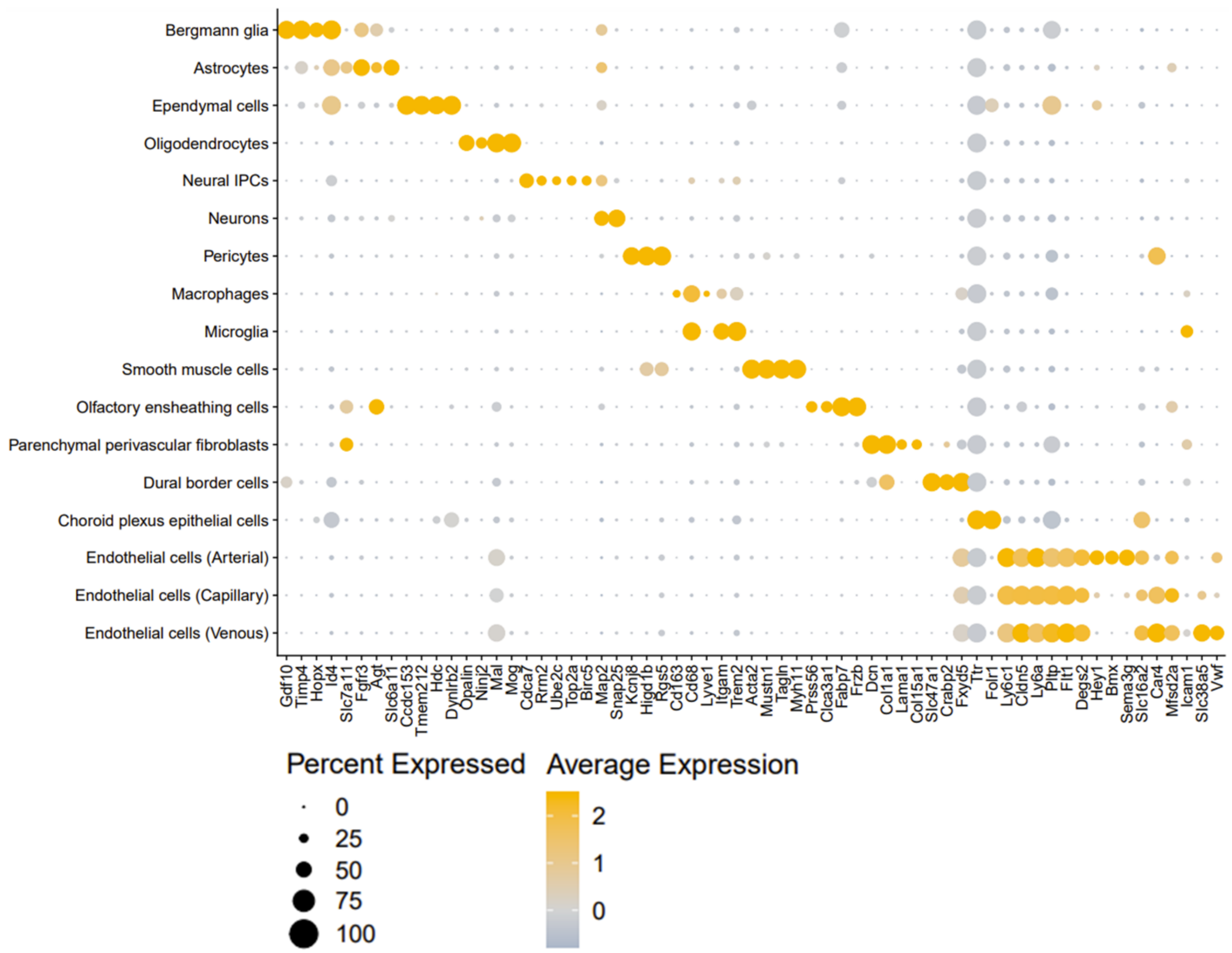


**Supplemental Figure 8: Quality control and annotations for scRNA-seq performed on whole brain**

A. Quality control metrics used for filtering in both samples.

B. UMAP with both samples shows minimal batch effects in cell-type clustering.

C. Smoothed ridgeline density plots of *Ppib* expression comparing expression between L2-siRNA$^{NTC}$ and L2-siRNA$^{PPIB}$ for all parenchymal populations not shown in Figure 4.

D. Dotplot showing markers used to identify cell types.

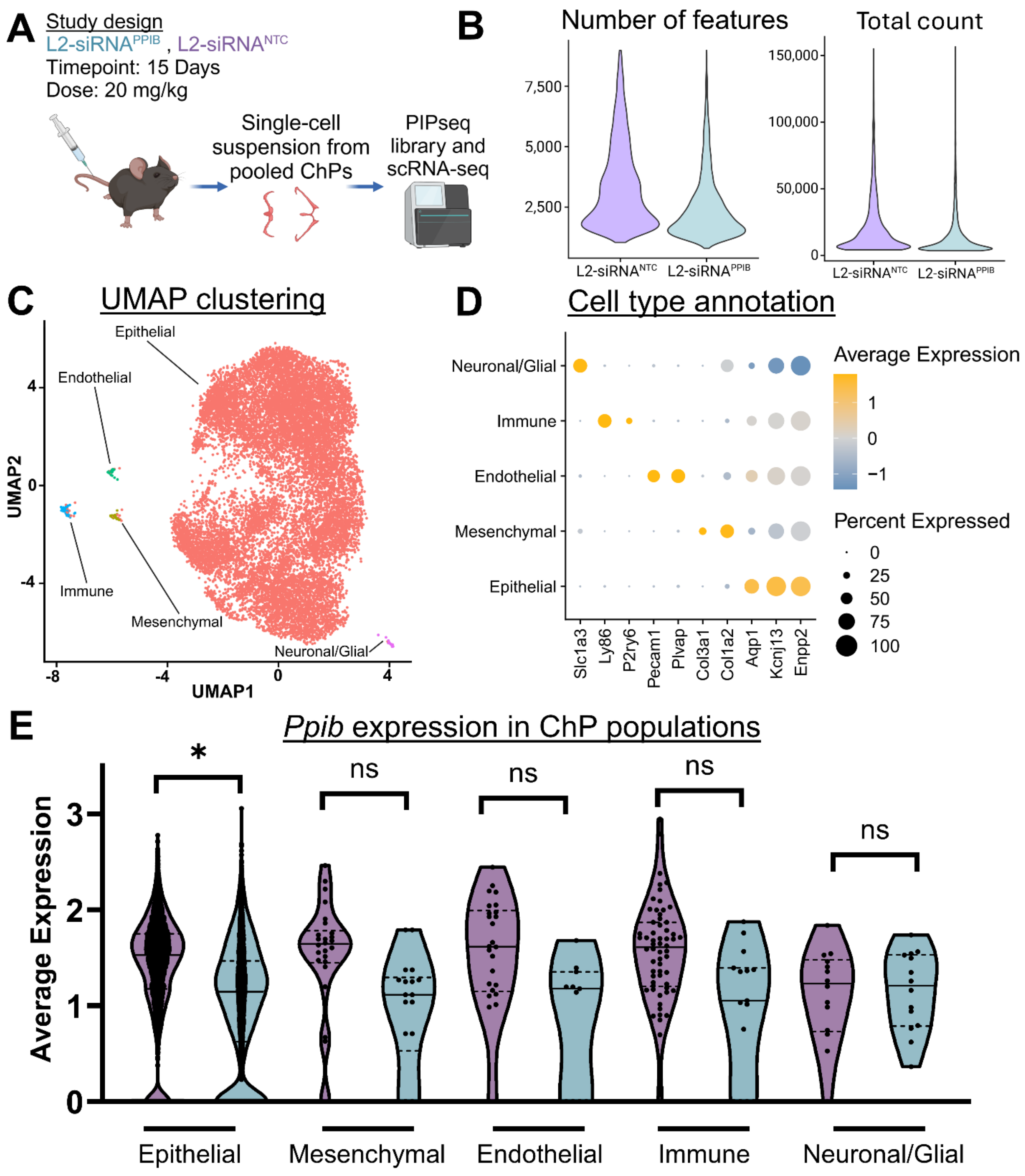


**Supplemental Figure 9: scRNA-seq of isolated choroid plexuses**

A. Experimental design to assess cell-specific gene silencing in the choroid plexus 15 days after 20 mg/kg injection, all choroid plexuses (lateral, 3rd, and 4th) were pooled from 4 mice and processed into a single cell suspension for sequencing.

B. Quality control metrics to filter out cells with low or high counts and features.

C. UMAP dimension reduction plot showing the five populations identified.

D. Dotplot showing markers used to identify cell types.

E. Violin plot showing changes in *Ppib* expression between L2-siRNA$^{NTC}$ and L2-siRNA$^{PPIB}$ for each cell type examined. Statistics were computed with a Wilcoxon test and Bonferroni correction for multiple comparisons.

| Name | Sequence (5'->3') | Location | Length | Reference |
|---|---|---|---|---|
| Ppib S | (MeA)*(fA)*(MeC)(fA)(MeG)(fC)(MeA)(fA)(MeA)(fU)(MeU)(fC)(MeC)(fA)(MeU)(fC)(MeG)(fU)*(MeG)*(fA) | 437 | 20 | Reynolds et al. 2004 |
| Ppib AS | VP(meU)*(fC)*(meA)(fC)(meG)(fA)(meU)(fG)(meG)(fA)(meA)(fU)(meU)(fU)(meG)(fC)(meU)(fG)*(meU)*(fU) | 437 | 20 | Reynolds et al. 2004 |
| Htt S | (MeU)*(fA)*(MeU)(fA)(MeU)(fC)(MeA)(fG)(MeU)(fA)(MeA)(fA)(MeG)(fA)(MeG)(fA)(MeU)(fU)*(MeA)*(fA)*(Cy5) | 10150 | 20 | Alterman et al. 2015 |
| Htt AS | VP(meU)*(fU)*(MeA)(fA)(MeU)(fC)(MeU)(fC)(MeU)(fU)(MeU)(fA)(MeC)(fU)(MeG)(fA)(MeU)(fA)*(MeU)*(fA) | 10150 | 20 | Alterman et al. 2015 |
| NTC S | (fU)*(MeU)*(fC)(MeU)(fC)(MeC)(fG)(MeA)(fA)(MeC)(fG)(MeU)(fG)(MeU)(fC)(MeA)(fC)*(MeG)*(fU) | NA | 19 | NA |
| NTC AS | VP(MeA)*(fC)*(MeG)(fU)(MeG)(fA)(MeC)(fA)(MeC)(fG)(MeU)(fU)(MeC)(fG)(MeG)(fA)(MeG)*(fA)*(MeA) | NA | 19 | NA |
| Ppib PNA probe | 5'/N Cy3-OO-AACAGCAAATTCCATCGTGA 3'/C | NA | 20 | Reynolds et al. 2004 |
| Key: S=sense strand, AS=anti-sense strand, *=phosphorothioate, Me=2'-O-Methyl, f=2'fluoro, VP=vinyl phosphonate, O = ethylene glycol linker | | | | |

**Supplemental Table 1: siRNA sequences used throughout manuscript.** Fluorescent biodistribution studies were performed with Cy5-labeled *Htt* sequences and knockdown studies were performed with *Ppib* and NTC sequences.
